# Social rank-order stability of mice revealed by a novel food competition paradigm in combination with available space competition paradigms

**DOI:** 10.1101/2024.10.15.618390

**Authors:** Meiqiu Liu, Yue Chen, Rongqing Chen

## Abstract

Psychological, behavioral and biological studies on social organization and competition with animal models are boosting. The mouse has been recognized as a valuable and economic model animal for biomedical research in social behaviors, however, currently available food competition paradigms for mice remain limited. Discrepant paradigms involving different competitive factors, such as physical strength vs psychological features, muscular confrontation vs threat perception, and boldness vs timidity, may produce task-specific win-or-lose outcomes and lead to inconsistent ranking results. Here, we developed a food competition apparatus for mouse, in which contenders were a pair of mice eager to take over the same food pellet placed under a movable block in the middle of a narrow chamber where they were separated to the either right or left side. This Food Pellet Competition Test (FPCT) was designed to (1) provide researchers with a choice of new food competition paradigm and (2) expose psychological factors influencing the establishment and/or expression social status in mice by avoiding direct physical competition between contenders. Meanwhile, we wanted to evaluate the consistency of social ranking results between FPCT and typically available space competition paradigms—tube test and warm spot test (WST). We hypothesized inconsistency of rankings of mice tested by FPCT, tube test and WST as they possess different targets for mice to compete and different factors determining competitiveness. Interestingly, application of FPCT in combination with tube test and WST discovered unexpected consistency of mouse social competitivity and rankings in a grouped male or female mice that were housed in an either 2-or 3-member cage, most likely indicating that the status sense of animals is part of a comprehensive identify of self-recognition of individuals in an established social colony. Furthermore, the FPCT may facilitate researches on social organization and competition, given its reliability, validity, and ease of use.

## Introduction

For gregarious animals and human beings, both competition and cooperation between individuals or populations are fundamental social behaviors important for the survival and evolution of the collective strains and species. Social competitions occur naturally to determine the ownership or priority of living territory, food, water, mates, other resources and non-resource requirements ^1,2^. Through these competitions, social hierarchies from the dominant to the subordinate are established within a living group, where individuals at higher social status are granted with corresponding priorities of resources and non-resource requirements like living space, food availability, reproduction and safeguard ^1,2^. The establishment of dominance hierarchies reduces the intensity and frequency of mutual aggression within groups and strains, so to maintain their inner stability and fulfil outer assignments ^3^. Impaired awareness of social competition has been documented in individuals with autism spectrum disorder (ASD) ^4,5^, and reduced social interaction has been characterized in corresponding animal models ^6^. Similarly, maladaptive responses to social status loss has been associated with patient depressive disorders ^7,8^ and animal models of depression ^9,10^.

Although humans and primates exhibit complex social interactions that are relatively easy to observe, their use for biological research is practically limited. In contrast, rodents —particularly mice and rats—have been emerging to serve as valuable and cost-effective models for studying the biological basis of social cognition, social interaction, and social organization ^9,11,12^. Among the social competing objects, resources of living space, water and food are fundamentally essential for the survival of animate beings. Therefore, living space, water and food competitions are most frequently used to investigate social competitive behaviors and hierarchical ranking of rats or mice ^2,13,14^. The tube test is a simple and reliable method for assessing social hierarchy by simulating the competition for living space among animals. In the tube test, the dominant mice largely consistently squeeze out the weaker ones from the narrow tube, indicative of an overwhelming space demand of mice at higher rank ^15^. Warm spot test (WST) is another space competition paradigm where a pair of mice compete for pitiful small warm corner in a freezing cage^12,16^. Some food competition tests have also been applied to animals such as rats ^17–19^, chickens ^20^, pigs ^21^, and mice ^22–25^. Larger body size is an advantage of rats, chickens and pigs that makes detecting behavioral patterns easier by the experimenters or monitoring video, but the food competition procedures for those animals are difficult to be applied to mice due to its much smaller size. Both mice and rats are rodents and mostly used experimental animals, but rats are more socially tolerant and less hierarchical than mice^26,27^.

Relative to space competition, food competition tests for mice have been designated and applied less commonly in animal studies despite its long history ^28–30^. Several issues could be thought to be the underlying limitations for the application of food competition paradigms. First, there are methodological issues in some of these approaches, such as long video recording duration and difficulty in analyzing animal’s behaviors during competitive physical interaction in videos, hindering their application by laboratories that cannot afford sophisticated equipment ^23,24,31–35^ and analysis ^36^. Second, aggressive behaviors often occur during competition in these approaches as the mice compete in a shared arena. Third, the prolonged food deprivation is usually required to increase the effortful food competition. For the second and third issues, on one hand, the aggressiveness/being aggressed and intense food deprivation cause stress responses and changes of physiological state of animals ^10,24,28,29^. On the other hand, competition in a same arena conflates multiple determinants contributing to the outcomes of competition involving physical aspects—such as muscular strength, vigorousness of fighting, bite wounding—and psychological aspects—such as boldness, focused motivation, active self-awareness of status. This feature makes interpretation of the outcomes of competition difficult to discriminate the dominant behaviors and aggressive behaviors. So far, the food competition tasks had not been conceived to separate the physical aspects from psychological aspects interpreting the mice’s winning/losing.

Therefore, to have a new choice of food competition paradigm for mice, and to facilitate the exposure of psychological aspects contributing to the winning/losing outcomes in competitions, we generated a convenient, easily operative and peaceful food competition paradigms for a pair of mice competing for a food pellet after mild calorie restriction. The FPCT was purposed for mice food completion without physical contact during competition, but tube test and WST were of space competition during which the mice need direct physical contact. Thus, we expected inconsistent evaluation results of competitiveness and rankings if we compared FPCT with typically available competition paradigms—tube test and WST.

## Results

### FPCT did not detect significant difference in the winning/losing results between unfamiliar non-cagemate male mice

The experimental procedure of FPCT consists of habituation, training and test (Figure 1A). After handling the mice in the homecage and experimental room, the mice were trained to enter the arena alternatively from the left and right gates to be familiar with the chamber arena (Figures 1B, S1 and S2), followed by next step to find a small food pellet in the trough in the middle of the chamber floor (Figures 1C and S2). At the end of this step of training, the mice would go directly for the food pellet when they entered the arena (Video S1). Then, a movable block was hung up at the roof of the chamber (Figures 1D, S1 and S2). The bottom of the high rectangular block was positioned nearly right above the trough so that the mice could not possess the food pellet unless they pushed the block away. The block is transparent and dug with holes at the lower part, enabling the mice see and smell the food pellet (Figure 1D). The mice were trained to learn to move away the block and reach the food pellet twice daily (Figure 1D). Usually, it took 4-5 days for the mice to learn to walk straightforwardly to the obstacle once they entered the chamber and move the block easily to expose the food pellet (Figure 1E and Video S2). Orientation of the entries did not generate bias of performance of mice in the apparatus (Figure 1E).

**Figure 1.**
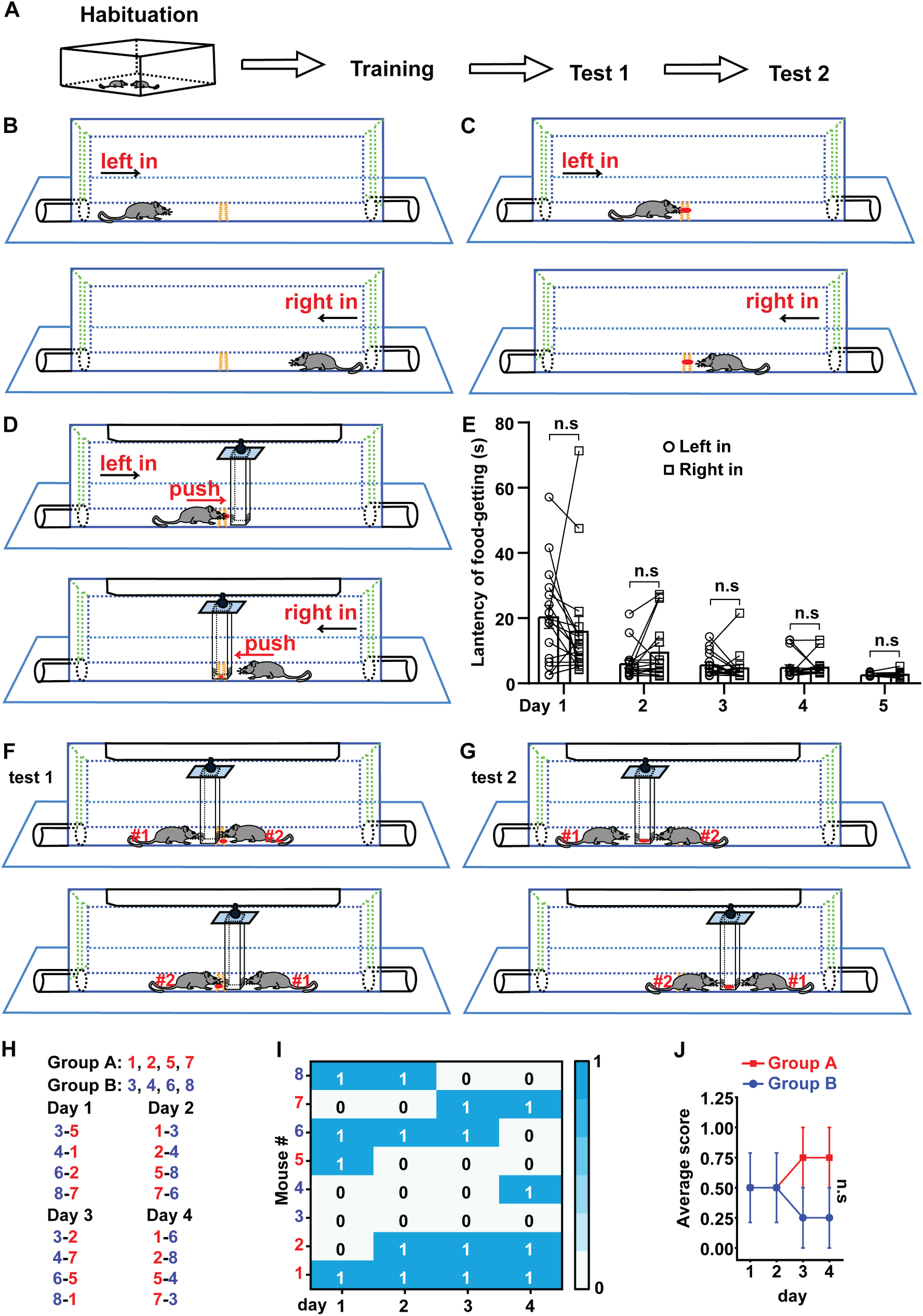
Schematic drawing of FPCT procedure and measuring the non-cagemate male mice based on the winning/losing outcomes of FPCT. (A) The overall procedure of FPCT experiment. (B) Habituation to the arena where the mice entered alternately from left and right sides. (C) Training to find food pellet without the existence of the movable block. (D) Training to get the food pellet placed under the movable block. (E) Statistical data of latency of food-getting (the time from entering the arena to eat the food) showing the progress of training in (D). paired Student’s t test, n = 16, n.s. stands for non-significant difference. (F) Direct win-lose test via analyzing food occupation (test 1). (G) Indirect win-lose test via analyzing mouse’s attempt to gain the food (test 2). (H) Eight single-housed mice were numbered and randomly divided into group A and B after being trained. One of the mice in group A matched only once with one of the mice in group B in the 4 days of test 1. The number listed on the left-right indicated the corresponding mice would enter the arena from left-right entry. Score 1 and 0 indicated winning and losing the food competition, respectively. (I) Heatmap showing the outcomes food competition of the non-cagemate male mice ranked with FPCT. (J) Average score of the two groups acquired in the competition. Unpaired Student’s t test = 4, n.s. stands for non-significant difference.

In the test section, two mice were allowed to enter the separated rooms of the chamber simultaneously from one of the two entries, competing for the food pellet either placed in the trough below the block (test 1, Figure 1F)—where the pallet was accessible if the block was pushed away—or on the inner bottom of the block (test 2, Figure 1G) —where the pallet was inaccessible. The two mice were able to see and sniff each other through the transparency and holes of the block, as well as the gap between the block and chamber.

In the first experiment utilizing the FPCT, we tested age-and weight-controlled unfamiliar non-cagemate male mice to examine whether there was obvious non-social priority-associated competitiveness expressed in the FPCT. Eight single-housed mice were trained and then randomly divided into groups A and B for competition test 1. Within the 4 consecutive days of competition test there were totally 16 trials, as one mouse in a group matched one of mice from the other group once daily and the matched two players participated in the contest only once in the 4-day schedule (Figure 1H). In each trial of the test 1, the mouse which obtained the food pellet was credited a score of 1 (time of winning), while the mouse which failed to obtain the food pellet was credited a score of 0 (Figure 1I). Statistical data found similar average scores acquired by the two groups of mice (Figure 1J), illustrating that no non-social priority-associated competition factor was obviously involved in the FPCT with the sex-, age-and weight-controlled unfamiliar mice.

### FPCT identified different social ranks between 2-cagemate male mice, verified by tube test

Social hierarchy or status of many animals is largely arranged by the outcomes of winning/losing social competitions, manifesting as dominant or subordinate behaviors in a well-established social colony. To see the efficacy of FPCT in detecting the social status of mice in a well-established social colony, we conducted 4-day of FPCT test 1 containing 1 trial per day for each of the 16 pairs of cagemate male mice after training (Figure 2A). In the neighboring 2 days, each mouse was allowed to enter the chamber from different entries. In each trial, the mice were scored 1 for winning and 0 for losing the food pellet. In the final trial, the mouse obtaining the food pellet was ranked #1 and considered as the final winner, while the mouse failing to get the food pellet in the match was rank #2 and claimed as the final loser. Retrograde analysis of the time spent to get the food after entering the arena gate in the last day of training showed that the winners and losers displayed same level of training to get the food and equivalent motivational craving for the food (Figure 2A). However, when there was competition in the test the winners tended to win the pellet overthought all the 4 trials (Figure 2A). Notably, the presence of an opponent during the test did not significantly change the latency to get the food, comparing with last day of training when there was no opponent (Figure 2A). In all the 16 pairs of mice, only 1 pair exhibited alternating winning/losing outcomes with an inter-trial consistency at 50% (Figure 2B), while all the other 15 pairs kept 100% consistency of winning/losing readouts (Figure 2C). Averagely, the competition results were stable throughout the 16 trials (Figure 2D) and a rate of consistency reached 96.88% (Figure 2E). In the test 2 (indirect win-lose test via analyzing mouse’s attempt to gain the food), we observed that the winners determined by the final trial of test 1 tried harder to get the food pellet by spending more time to push the block within the single 2-minute test, a limited time to avoid distinction of attempt (Figure 2F). Thus, these data demonstrate that FPCT are effective to rank the established social status of mice, and that consecutive 4 trials (one trial daily) might be least required for the FPCT to achieve readout consistency and reliability.

**Figure 2.**
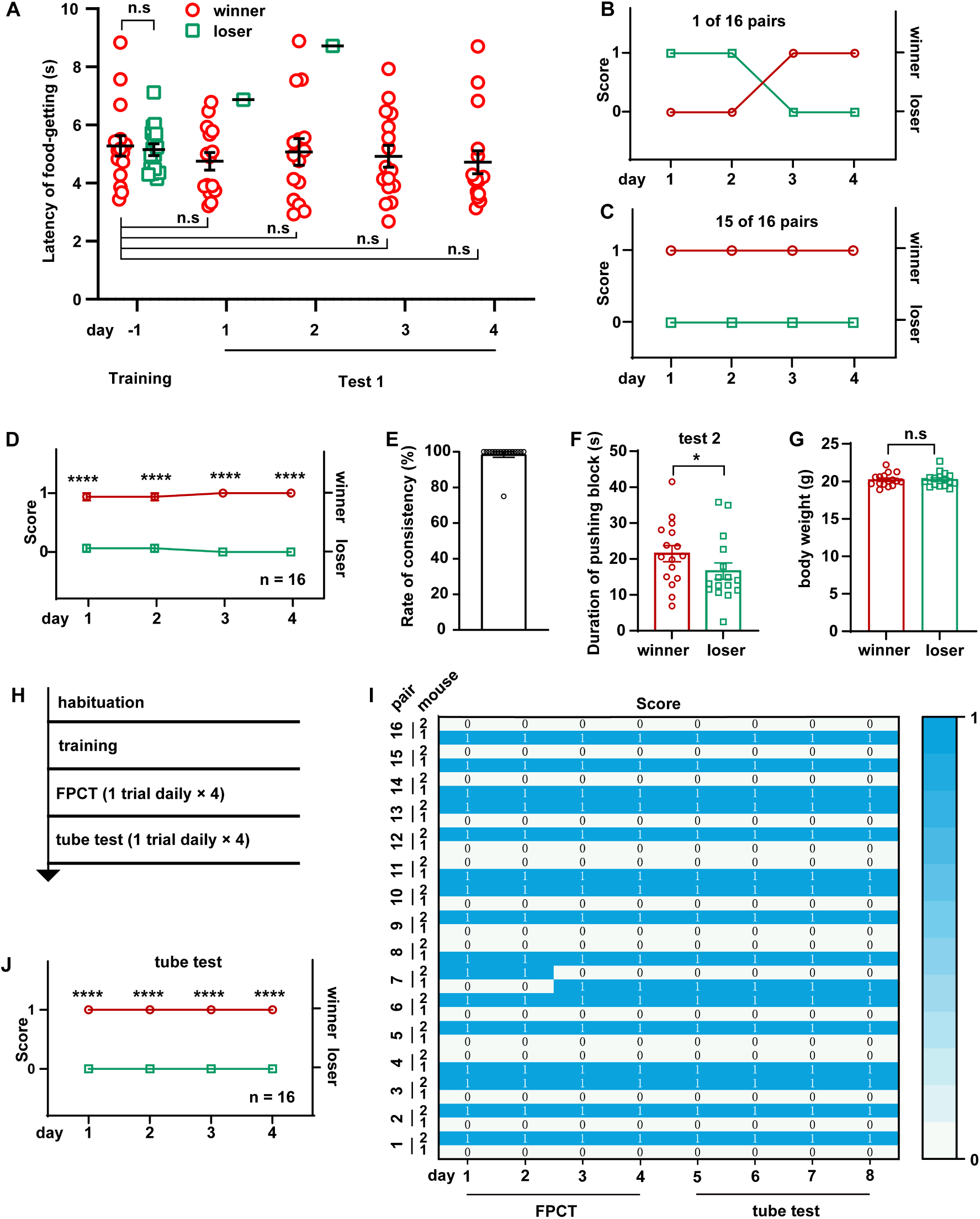
Ranking the 2-cagemate male mice with FPCT and tube test. (A) Statistical data showing the latency of food-getting from the last day of training until the end of the test 1. Two-way ANOVA multiple comparisons, n = 16 pairs, n.s. stands for non-significant difference. (B) 1 of the 16 pairs of cagemate male mice exhibited alternating winning/losing outcome in FPCT test 1. (C) 15 of the 16 pairs of male mice exhibited fully congruent winning/losing outcome in FPCT test 1. (D) Statistics of ranking of male mice over through 4 trials (1 trial daily) of FPCT test 1. Two-way ANOVA, n = 16, \*\*\*\**P* < 0.0001. (E) Average rate of consistency within trials. For each pair, rate of consistency was calculated as the percentage of the number of trials (in all 4 trials) resulting in the outcome same as the 4^th^ trial (n = 16). (F) Statistics of duration spent on pushing block in FPCT test 2. The winner or loser identity was determined by the last trial of test 1. Paired Student’s t test, \**P* < 0.05, n = 16. (G) Body weight of male mice measured after FPCT. Unpaired Student’s t test, n.s. stands for non-significant difference, n = 16. (H) Timeline of experiments showing tube test was conducted after FPCT. (I) Heatmap showing the outcomes of social competition of paired male mice ranked with FPCT and tube test. (J) Statistics of ranking of male mice over through 4 trials (1 trial daily) of tube test. Two-way ANOVA, n = 16, \*\*\*\**P* < 0.0001.

Many factors may contribute to the competitive outcomes in mice, such as age, sex, training level, physical strength, and intensity of psychological factors. We were interesting to understand which factor determines the outcomes of mice’s winning/losing in the FPCT match. First, effects of age and sex on wining/losing outcomes were excluded as the mice were age-matched and solely males. Second, the weight of mice was also excluded to determine the competency possibly by providing physical strength of pushing, as the weight difference between paired contender mice before training was controlled in less than 10%, and no difference between the winners and losers in weight measured again after finishing the food competition tests (Figure 2G). Third, biased training level should produce distinct likelihood of winning and losing, but retrospective comparison of the duration they took to obtained the food pellets in the last day of training when there was no competition reported no difference in the latency to get the food pellet between the winners and losers (Figure 2A), indicating that the mice were equally trained and motivated. Forth, the major physical contribution to the competition based on violence aggressiveness and fighting strength were minimized in this competitive scenario due to separation of the mice by the obstacle. Thus, we postulated that the winning/losing outcomes in this paradigm mainly rely on mice’s state of psychological factors, for example, intensity of motivation, self-awareness of social status, memory of winning/losing experience in the homecage, fear of revenge when returning the living cage. Most possibly, self-awareness of social status plays a determinant role in the outcomes.

To see whether social ranking revealed by the FPCT is consistent with the tube test that represents the simple and robust behavioral assay for space competition ^15^, we continued the experiments to rank the mice using tube test following FPCT (Figures 2H and 2I). We found that consistency of outcomes between the FPCT and tube test were 100 percent, as the winners and losers in the last trial of the FPCT continued consecutively to be the winners and losers, respectively, during the 4 consecutive trials of tube tests (Figures 2I and 2J). Therefore, the rank-order examined by with the FPCT is fully verified by the tube test.

### Social ranking of 2-cagemate female mice using FPCT, tube test and warm spot test

It is widely accepted that males, including human beings and animals, are evolutionarily more eager to be dominant, more aggressive and more hierarchical, but it is arguable regarding whether females have less competition and looser social organization ^37–39^. On the base of investigation of aggressive behaviors, it has been reported that the social colony of female mice is also hierarchical ^30,40^. Taking use of this newly designed mouse competition task in which dependence of competency on physical aggressiveness is minimized, we examined whether intra-female social status between two cagemate female mice was observable in the FPCT context. Notably, the FPCT test 1 showed that 3 in 10 pairs of cagemate female mice exhibited alternating winning/losing outcomes, while the majority of cagemates (7 in 10 pairs) showed fully congruent grades (Figures 3A-3E). Averagely, the 10 female pairs displayed stable rankings over through all trials (Figure 3F). The overall rate of inter-trial consistency was 90% (Figure 3G), which was not significantly different from male mice (*P* > 0.05 comparing data in the Figures 2E and 3G by Mann-Whitney U test). In the test 2, we observed that winner mice in the test 1 paid more efforts trying to get the food pellet (Figure 3H). The body wights examined after the tests (Figures 3I) were similar between the winners and losers, so the two parameters were not likely to contribute to the competency. Interestingly, retrospect analysis of training data displayed similar training level of food-getting and craving state for food (Figures 3A). However, the presence of an opponent during the test 1 could significantly disturb the female mice and increase the its latency to get the food, comparing with last day of training when there was no opponent (Figure 3A). These FPCT results suggest that the social relationship of female mice is also stratified.

**Figure 3.**
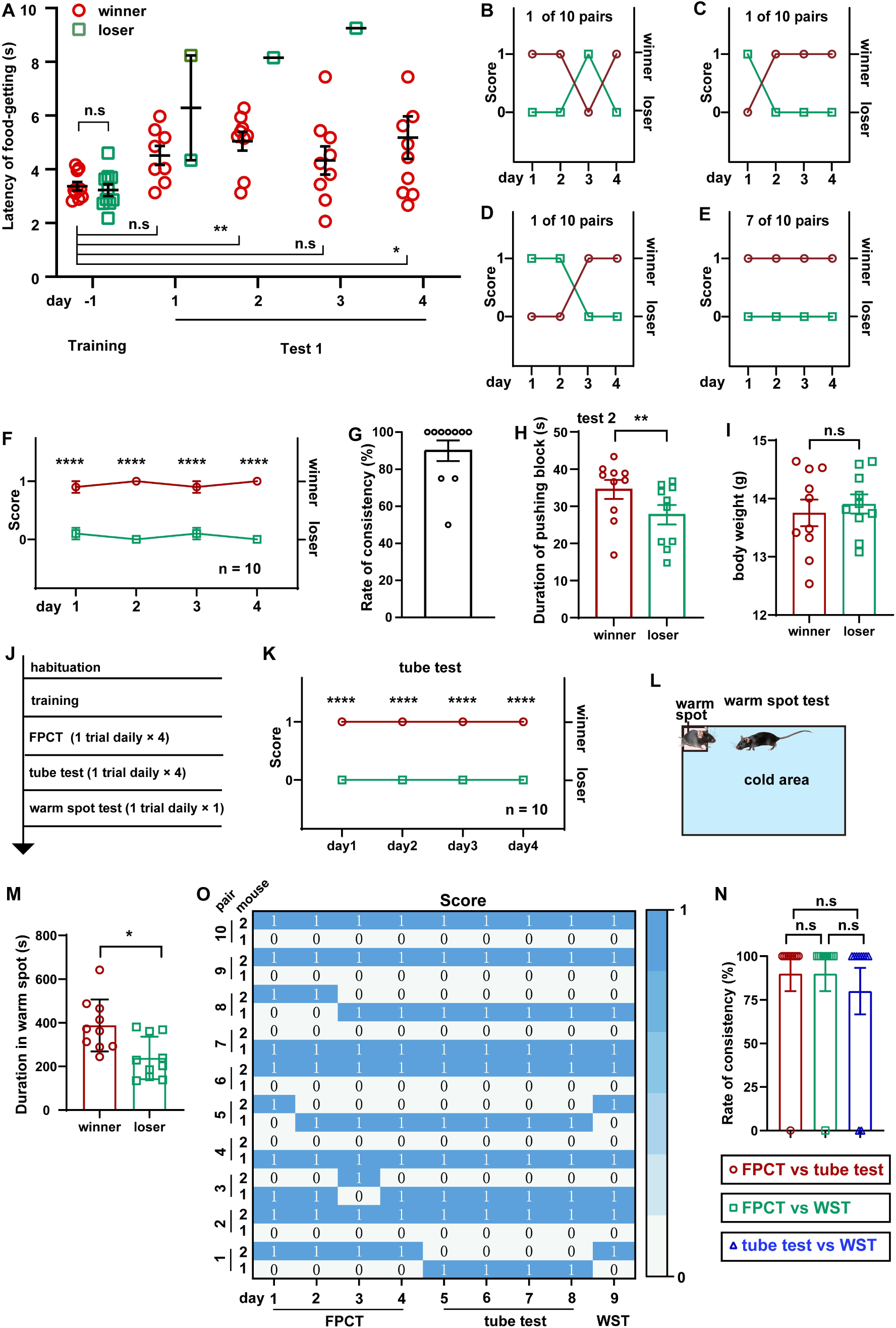
Ranking the 2-cagemate female mice using FPCT, tube test and warm spot test (WST). (A) Statistical data showing the latency of food-getting from the last day of training until the end of the test 1. Two-way ANOVA multiple comparisons, n = 10 pairs; \**P* < 0.05, \*\**P* < 0.01, n.s. stands for non-significant difference. (B-D) 3 of the 10 pairs of cagemate female mice exhibited alternating winning/losing outcome in FPCT test 1. (E) 7 of the 10 pairs of female mice exhibited fully congruent winning/losing outcome in FPCT test 1. (F) Statistics of ranking of female mice over through 4 trials (1 trial daily) of FPCT test 1. Two-way ANOVA, n = 10, \*\*\*\**P* < 0.0001. (G) Average rate of consistency within trials. For each pair, rate of consistency was calculated as the percentage of the number of trials (in all 4 trials) resulting the outcome same as 4^th^ trial (n = 10). (H) Statistics of duration spent on pushing block in FPCT test 2. The winner or loser identity was determined by the last trial of test 1. Paired Student’s t test, \**P* < 0.05, n = 10. (I) Body weight of female mice measured after FPCT. Unpaired Student’s t test, n.s. stands for non-significant difference, n = 10. (J) Timeline of experiments showing tube test and warm spot test were conducted after FPCT. (K) Statistics of ranking of female mice over through 4 trials (1 trial daily) of tube test. Two-way ANOVA, n = 10, \*\*\*\**P* < 0.0001. (L) Schematic of the warm spot test. (M) Cumulative duration of mice occupying the warm spot. The winner or loser identity was determined by the last trial of FPCT. Paired Student’s t test, n = 10, \**P* < 0.05. (O) Heatmap showing the outcomes of social competition of paired female mice ranked with FPCT, tube test and WST. (P) Rate of consistency between FPCT and tube test (day 4 vs day 5), FPCT and WST (day 4 vs day 9), as well as tube test and WST (day 8 vs day 9). One-way ANOVA test, n = 10, n.s. stands for non-significant difference.

In addition to the tube test, WST, the regional urine marker and the courtship ultrasound vocalization test are established behavioral methods for determining social rank in mice ^12,14,16,17,32,41–46^. Tested animals in different tasks may pursue specific goals, and utilize their specific expertise, strength and skills. Thus, we assumed that different competition paradigms may produce inconsistent social ranking readouts of the same social colony, resulting from various types and levels of competition. To check this assumption, we continued to assess these female mice using tube test and WST following FPCT (Figure 3J). As expected, some of the trials exhibited inconsistent rankings the three tests. However, most of the trials unexpectedly showed that mice maintained the winner or loser identity acquired in the FPCT in subsequent tube test and WST (Figures 3K-3M). Overall, no difference was observed in the rate of inter-trial consistency comparing FPCT with tube test, FPCT with WST, and tube test with WST (Figure 3N). These data illustrate that mouse social competency and status of established female mice colony are overall stable so that a well competitive subject exhibits dominance not just in a specific, but in a variety of contexts, despite that the different contexts contain distinct competitive factors like food and space, or competitive aspects like physical strength and psychological factors.

### Social ranking of triads of male mice using FPCT, tube test and WST

Social organization of the bigger crowd could be much more complicated than the simplest 2-member group. It is controversial whether the dominate and subordinate roles in a larger group of mice tend to be mobile or rigorous in a variety of contexts ^35,47,48^. To probe this issue, we raised 3 male mice in a cage and then following adequate habituation and training we ranked their hierarchies using FPCT, tube test and warm spot test sequentially (Figure 4A). In the FPCT and tube test, the 3 cagemate mice contested in a round-robin style as the both were designated for two contenders in a time. The ranking result showed that 6 in 9 groups of mice displayed some extent of flipped ranking (Figures 4B-4G), and only 3 in 9 groups displayed continuously unaltered ranking (Figure 4H). Averagely, in the totally 27 trials consisting of 12 trials of FPCT, 12 trials of tube test and 3 trials of WST, an obvious stable linear intragroup hierarchy was observed across all the trials and tasks (Figures 4I and 4J). Comparison of inter-task consistency revealed that the ranks assessed by FPCT, tube test and WST did not differ from each other (FPCT vs tube test, 85.19 + 17.95%; FPCT vs WST, 81.59 ± 17.95%; tube test vs WST, 81.48 ± 17.95%; *P* > 0.05, one-way ANOVA test. Figure 4K). These results, together with the results of female mice tested by FPCT, tube test and WST, illustrate that the dominate and subordinate roles in an established group of mice are rigorously linearly stratified so that the status can be manifested in various contexts.

**Figure 4.**
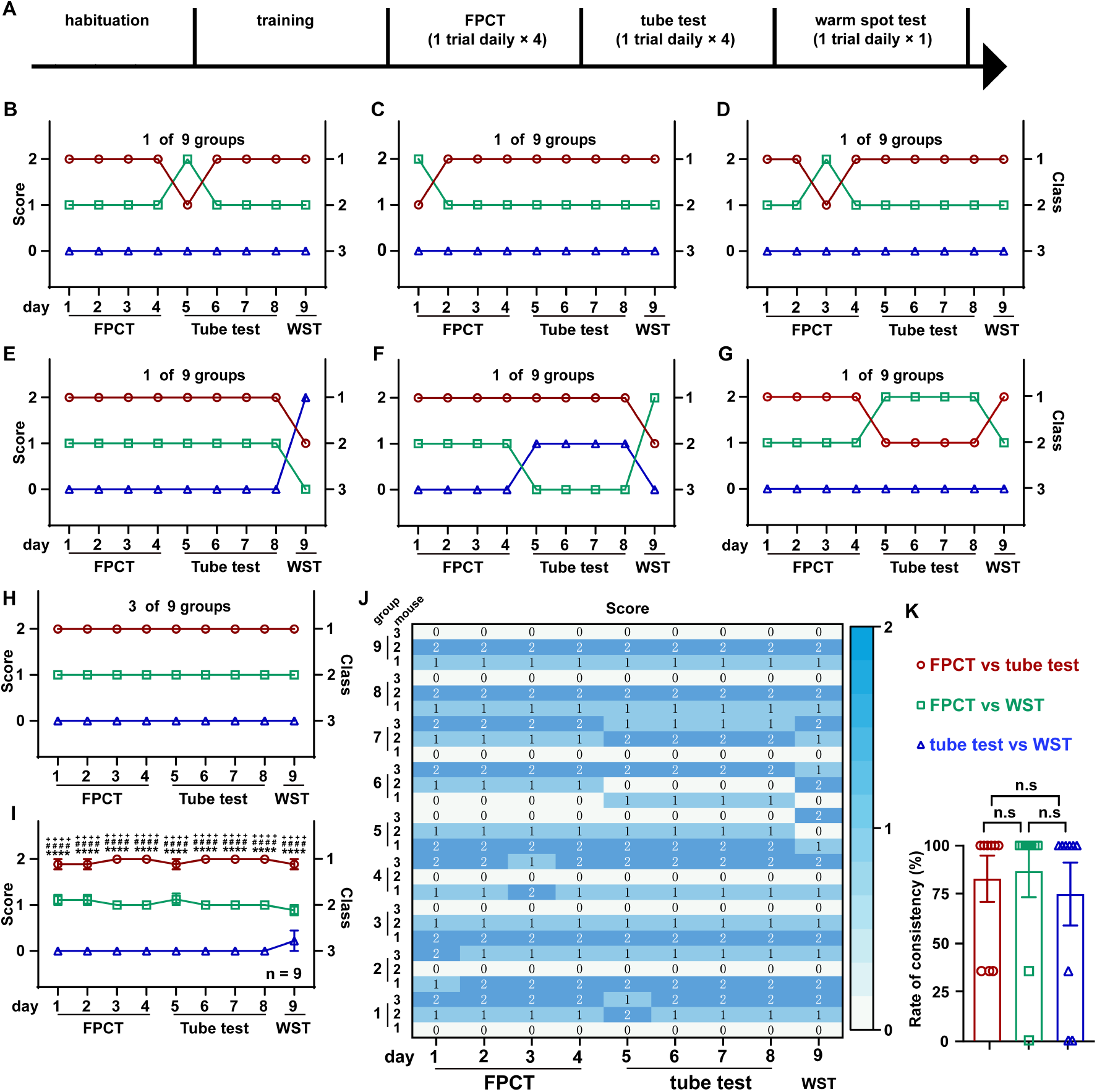
Ranking the triads of male mice using FPCT, tube test and WST. (A) Timeline of experiments to rank 3-cagemate male mice using FPCT, tube test and WST sequentially. In the FPCT and tube test, the mice contested in a round-robin style within the each 3-cagemate group. (B-G) 7 of the 10 groups of male mice exhibited alternating winning/losing outcome during the whole competition tasks. (H) 3 of the 10 groups of male mice exhibited fully congruent winning/losing outcome during the whole competition tasks. (I) Statistics of ranking of mice over through the whole competition tasks. Two-way ANOVA test, n = 10, ^++++^*P* < 0.0001 comparing FPCT and tube test, \*\*\*\**P* < 0.0001 comparing FPCT and tube test, ^####^*P* < 0.0001 comparing tube test and WST. (J) Rate of consistency between FPCT and tube test (day 4 vs day 5), FPCT and WST (day 4 vs day 9), as well as tube test and WST (day 8 vs day 9). One-way ANOVA test, n = 10, n.s. stands for non-significant difference. (K) Heatmap showing the ranking outcomes of grouped male mice during the whole competition tasks.

## Discussion

Social hierarchy is an innate feature of gregarious animals, usually established through social competition. Among the diverse contents for social competition, resources of living space and food are fundamentally critical priority for animals ^2,13^. Mice have been designated to participate in food competition tests for a long time, since at least the 1950s ^28,29^. In these applied food competition tests, social ranks were scored by calculating the total amount of food consumed by each mouse competing in the same chamber or cage, and/or analyzing their physical aggressiveness ^22,23,30,48^. Some common limitations for the wide distribution of these food competition tests are significant, including the occurrence of aggressive behaviors, requirement of prolonged food deprivation, calculation formulas, long video recording duration, and difficulty in discriminating individual animal’s behaviors during interacting with group members in videos ^23,24,31–35^. Food shortage serves as a strong driver to induce aggressiveness and competitive behaviors ^49^. However, in rodents, being physically aggressed and undergoing intense food deprivation can influence social competitive behaviors via disturbing the internal state ^24,31,50^ and triggering robust stress responses ^49–53^,.

Here, we developed and validated a food competition assay—FPCT, a simple and effective tool designed to provide a means of assessing social status of mice, favoring the growing scientific interest in understanding the neurobiological insights of social hierarchy. The potential influence of food restriction on competitive behavior was minimized in our task where the mice underwent only a 24-hour food deprivation period at the beginning of training, followed by restrict food supply to meet basic energy requirement. The aggressive situations, at least direct physical fighting, was prevented during this ranking test due to the separation of the mice in either side of the arena. The FPCT paradigm is conveniently and generally applicable for its simple procedure. First, the mice were housed together to habituate each other and establish hierarchical social structure. Then, the mice were trained individually to accommodate the chamber arena, know the existence and position of the food pellet, and learn to push away the block above the food pellet to get the food. Importantly to note, stable social hierarchy relies on adequate habituation and reliable ranking test is based on well training. Finally, the mice underwent the ranking test in the chamber, where they were allowed to enter from either the right or the left side of the chamber separated by the movable block to compete for the same food pellet under the block.

Concerning the interpretation of the ranking results, both static vs dynamic and linear vs despotic hierarchy organizations of grouped mice that have been documented ^35,47,48^. The diversity of ranking results might be attributed to various factors. For example, different paradigms containing different tasks that may enable different mice to express its proficient competitiveness such as physical advantage, psychological strength, and learning ability. Bodily conflict between contender mice in a shared space could make muscular strength be competitive. It is not sure if body size/weight affect social hierarchy as both correlation^35^ and non-correlation ^10,48^ between weight and hierarchy have been reported. Strategic task possibly promotes a smarter mouse to be a winner in the test even if it may in a subordinate role at homecage. Besides, group size of experimental animals ^54,55^, duration of habituation of group members ^55^, training level and test protocols (e.g., short-term vs long-term test) ^47^ should be considered as common points give rise to instability of the formation or expression of social hierarchy. In this novel food competition paradigm, we tested pairs and triads of same-sex mice with similar age and body weight, housed in the same cage for at least 3 days. We did not find significant difference in food priority within pairs of non-cagemates, whereas we detected overall stable winning/losing outcomes of cagemate pairs and triads. Considering that direct bodily competition between mice was avoided in the FPCT by separating the matching pair in either side of the chamber, the stable ranking results of the matches should be mainly determined by the psychological aspects of the contenders. Most possibly, it is a reflection of mouse’s self-awareness of social status.

Combining FPCT with typically available paradigms—tube test and WST, we addressed 3 questions: whether female mice are stably socially structured, whether FPCT is effective to discover hierarchy rankings across multiple cagemates, and whether different social tasks result in discrepant hierarchy rankings. First, we found that the majority of female pairs displayed overall stable hierarchy ranks, consistent with previous findings ^40,56^. We also detected 3 in 10 female cagemate pairs exhibiting fluctuant rankings during the 4-day FPCT test, but it was not significant from 1 in 16 male pairs. It has been revealed by agonistic behaviors in previous work that female social hierarchies are not as steep and despotic as male hierarchies ^39^. The discrepancy between this work and ours might be attributed to difference in mouse trains, group sizes, housing models and detecting methods. Second, by means of round-robin tests of multiple cagemates, both FPCT and tube test detected averagely linear intragroup hierarchies, consistent with each other. Third, considering that diverse competitive factors lying in different tasks. the ranking readouts of male pairs and triads examined by FPCT, tube test and WST were highly consistent with each other, revealing social rank-order stability of mice across the three types of examining tasks. Notably, the fluctuance of rankings occurs occasionally, suggesting that the social status of mouse could be dynamic and transitive in some occasions ^47^. It would be interesting to investigate whether the social hierarchies in larger sizes of mice group, especially female mice, are more dynamic and transitive. Alternately, the occasional intra-trial and inter-task ranking fluctuations might be a result of incompetency of paradigms to reflect the real intra-colony structure of mice.

In conclusion, our data suggest that hierarchical sense of animals might be part of a comprehensive identify of self-recognition of individuals within an established society. Hopefully, the FPCT will facilitates future studies to reveal more detailed properties of social organization, social competition and their underlying neurobiological mechanisms.

## Methods and materials

### Animals

All animal experiments were conducted according to the Regulations for the Administration of Affairs Concerning Experimental Animals (China) and were approved by the Southern Medical University Animal Ethics Committee. The mice used in this study were 6-8 weeks old C57/BL6J purchased from SPF (Beijing) Biotechnology Company (Beijing, China) and were raised in the environment with the relative humidity 50-75%, temperature 22-24°C and 12-12-h light-dark cycle. The mice were allowed to freely access to food and water unless undergoing food restriction requirement during experiments.

### Food pellet competition test (FPCT) setup

The experimental setup consists of a food competition arena (Figure S1) and a camera. The arena includes a base, a chamber, two entries, two doors, a movable block, a pulley system, a food trough and the roof (Figure S2) made up of a batch of wheels and a wheel track for the pulley system, as well as 15 pieces of acrylic plates for other parts listed with dimensions in Figure S2F. The movable block hung up under the track section separates the arena into the left and right compartments. It is necessary to apply some paraffin oil to the track to reduce the friction between the pulley and the track (Figure S3). The food trough is positioned in the middle of the chamber floor and nearly under the movable block unless the block is pushed away by mice. Food pellets are tiny delicious milk crackers. Mouse behaviors are monitored by a side view cameral with PlotPlayer software.

### FPCT procedure

The experimental procedure consists of habituation, training and test (Figure 1A). The details of the procedure are described below.

### Habituation

Contender pair of C57BL/6JNifdc mice with similar age and weight (the weight difference was less than 10%) were housed in the same cage for at least 3 days under a 12-12-h light-dark cycle. Before the formal training of mice, each mouse was handled by experimenters for 3-5 minutes per day for approximately 3 days. Overall, the habituation period lasts about 6-8 days.

### Training

Step 1, food restriction. Food restriction includes food deprivation and food limitation. First, the mice were deprived of food for 24 hours while water consumption remained normally to enhance the appeal of the food pellet to the mice. Then, after 24 hours of food deprivation, each cage of mice was given 10 g of food every morning to meet their daily food requirements until the end of the test.

Step 2, contextual familiarization (Figure 1B). At each time after mice homecage was translocated from animal facility to behavioral room, the cage lid was removed to allow the mice to freely explore the cage for about 1-2 minutes to become accustomed to the open roof. Before training to find the food pellet (step 3), each mouse was allowed to enter the arena chamber from the left side where the mouse was kept for 3 minutes before been gently driven out from the right side of the chamber to be back to homecage to have a 2-minute rest. Then, the mouse would enter the arena chamber from the right side and 3 minutes later was gently driven out from the left side. Arena familiarization was conducted 1 round per day for 2-3 days.

Step 3, training to find the food pellet (Figure 1C). The experimenter placed a small food pellet in the trough in the middle of the floor of chamber. Let one mouse enter the chamber from the left side and stay there until it found and ate the pellet. After that, the mouse was gently driven out of the chamber from the right side to have a 2-minute rest in its homecage. Then, the mouse would enter the chamber from the right side and stay there until it found and ate food. The training was repeated 1 round daily until each mouse directly went to the trough and ate the small pellet after entering the chamber (Video S1).

Step 4, learning to get the food pellet (Figure 1D). In this step, the small food pellet in the trough was nearly under the vertical block. As the block was transparent and had holes at lower portions facing to the entries, and it was movable thanks to the sliding wheels and the track at the top of the block attached to the roof, the mouse was able to learn to push away the block to get the pellet. The roof attached with track and block was lifted up after the mouse ate the pellet to allow the mouse go out of the chamber from the side opposite to the entry side. Each mouse was repeatedly trained to get the food pellet alternatively from left and right entry until they were able to harvest the pellet directly and easily without competition (Video S2).

### Test

Test 1, direct win-lose test via analyzing food occupation (Figure 1F). Test 1 required several trials, in the first of which the paired contenders entered the chamber from the opposite sides simultaneously. The mice entered the chamber from left or right entry alternatively in two consecutive trials. As there was only one small pellet placed under the movable block in each trial, the mouse that obtained the pellet was deemed as winner, while the other one was the loser (Video S3). To calculate the latency of food harvesting of mice, a starting state was marked when its four legs just entered the chamber arena (Figure S4). The time spent from starting state to food-getting for a mouse was calculated as latency of food harvesting. One competition test comprised at least 4 consecutive trials.

Test 2, indirect win-lose test via analyzing mouse’s attempt to gain the food (Figure 1G). In this test, a slightly larger pellet was placed in the inner bottom of the block, rather than in the trough under the block, so that the mice could see, smell but not access it. Once the mice entered the chamber, they would push the block and attempted to gain the food. The winner or loser was determined by times of pushing the block in 2 minutes (Video S4).

### Tube test

The detailed steps of tube test were described previously by Fan et al. ^15^. It consists of the following three steps: adaptation, training and test. The adaptation phase had been completed at FPCT, referred to as the habituation step of FPCT. In the training phase, the tail of the mouse was gently lifted and placed at one end of the tube, and when the mouse entered the tube, the tail was released and the mouse was allowed to pass through the tube. A plastic rod was used, when the mouse retreated or stagnated for a long time, to gently touched the tail so that the mouse could continue to move until it passed through the tube. The test phase required four consecutive days, and the social rank of the mice was ranked for four consecutive days. During the experiment, the camera was placed directly in front of the tube, and the whole process of the mouse experiment could be recorded.

### Warm spot test (WST)

The experimental apparatus utilized for the WST consisted of a rectangular behavior box with dimensions of 28 cm in length, 20 cm in width, and 40 cm in height. Prior to the commencement of the experiment, the bottom of the box was placed on ice to ensure that its temperature was approximately 0°C. A heating sheet measuring 2 cm by 2 cm was positioned in one of the inner corners of the box, providing sufficient space for only one mouse. The temperature of the heating sheet was maintained at 34°C, and this temperature was monitored using a temperature measuring gun. During the experiment, three mice were introduced into the box simultaneously, allowing them to move freely for a duration of 15 minutes. The entire experimental procedure was recorded via video, and the time each mouse spent on the heating sheet was subsequently quantified from the recorded footage.

## Statistical analysis

The experimental data were statistically analyzed using *Prism* 9 or origin 9.0 software and presented as mean ± SEM. Two-way or One-way ANOVA, unpaired or paired Student’s t test and Mann-Whitney U test were used to compare the difference between groups. Statistical significance level (*P* value) was set at 0.05.

## Author contributions

M.L. and Y.C. acquired the experimental data. M.L. analyzed the data. M.L. and R.C. established methodology and wrote the manuscript. R.C. conceived the project and supervised the research. All authors contributed to the finalization and approved the content of the manuscript.

## Funding

This work was supported by the National Natural Science Foundation of China (NSFC) (82271859 and 31771127) and the grant from the Natural Science Foundation of Guangdong Province (2023A1515010639).

## Availability of data and materials

The data and supplementary information supporting the findings of this study are within the paper and other information helping to understand this study could be available upon request from the corresponding authors.

## Declaration of interests

The authors declare no competing interests.

## Supplementary figure legends

**Supplementary figure 1 (Figure S1).**
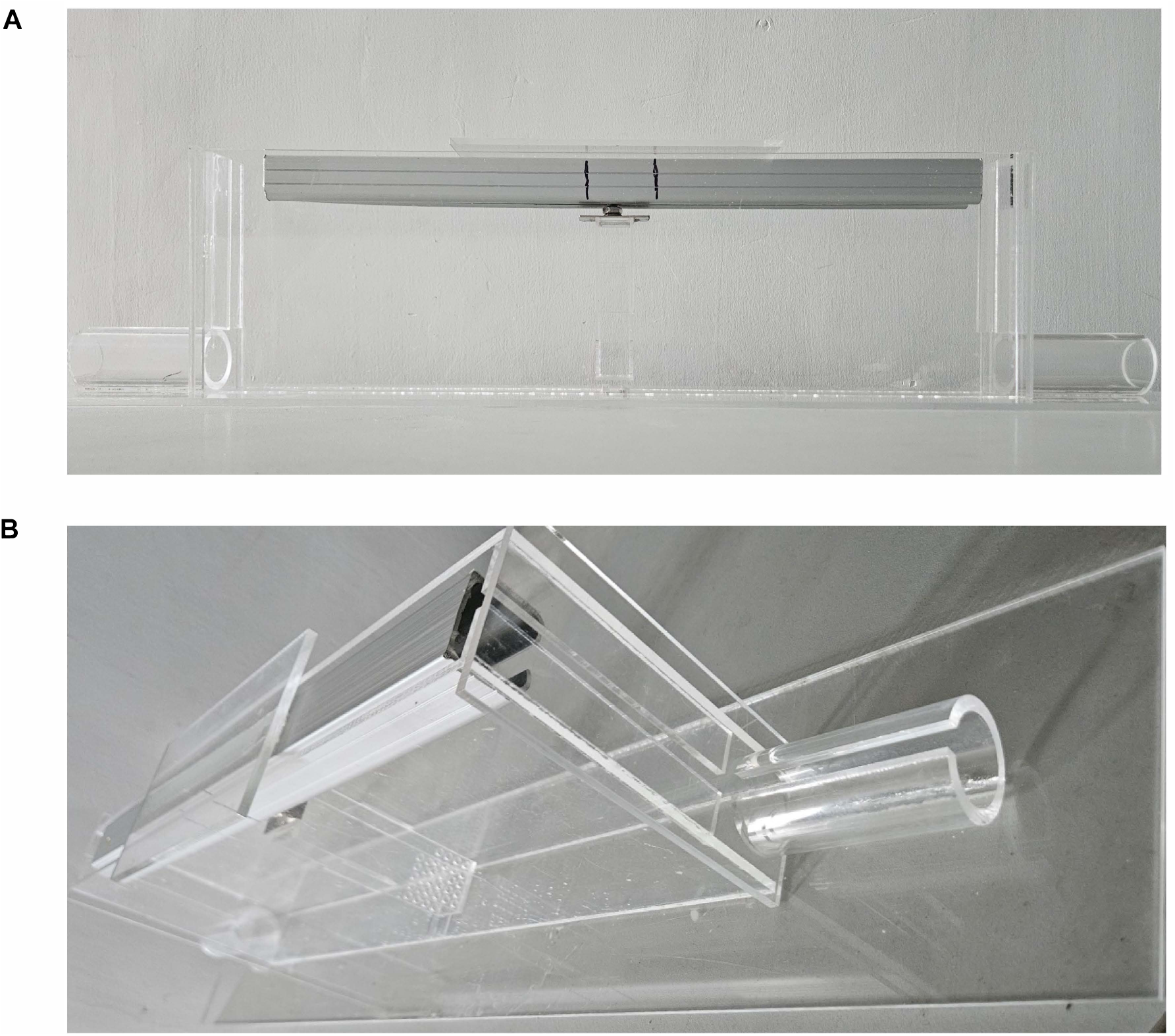
Photographs of food pellet competition test (FPCT) setup. (A) Frontal view of FPCT setup. (B) Side view of FPCT setup.

**Supplementary figure 2 (Figure S2).**
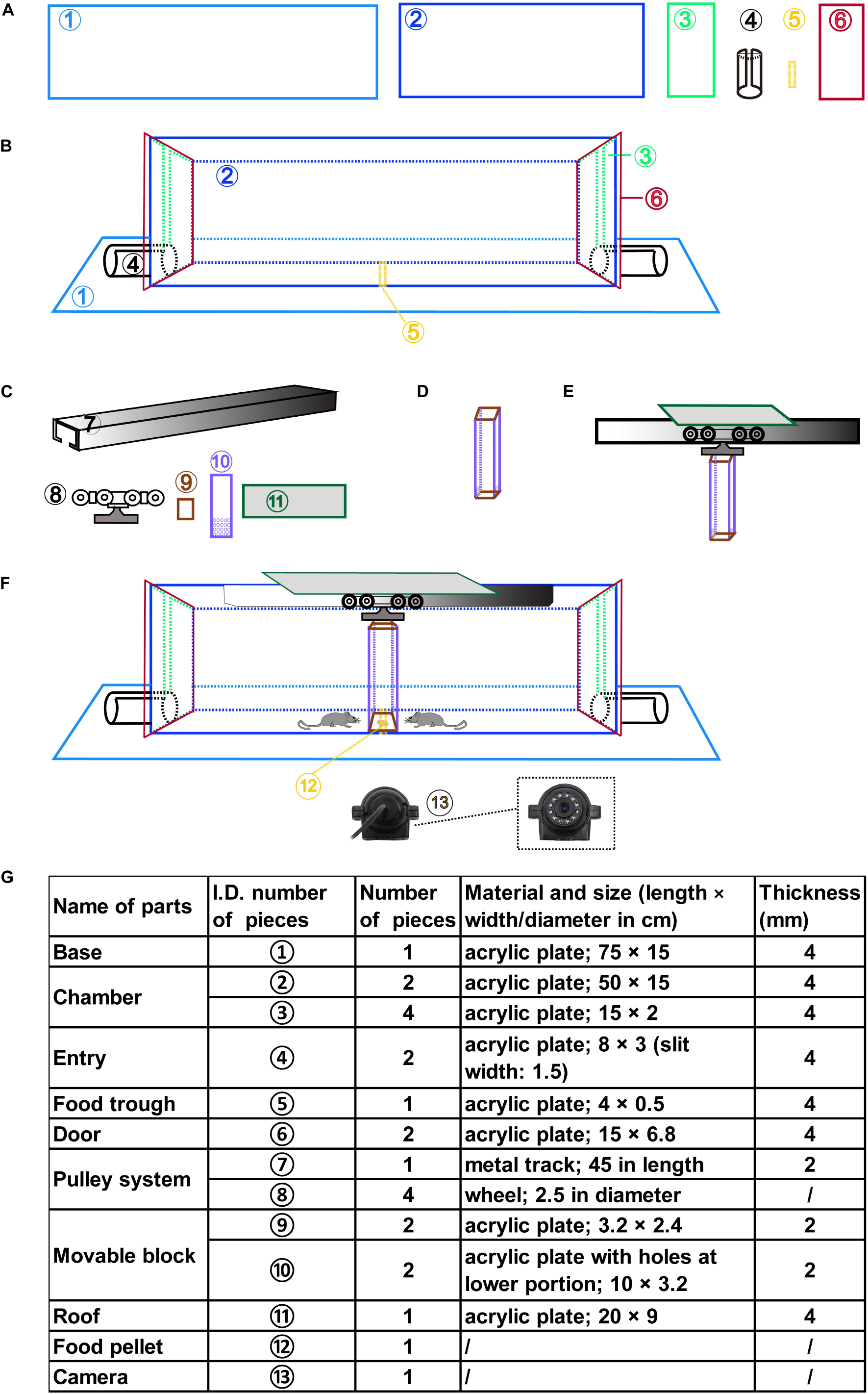
Schematic illustration of the assembling of the FPCT setup. (A-B) Depiction of the work pieces (A) to make up the main body of the arena (B). (C-E) Depiction of the work pieces (C) to make up the movable block (D) attached to the pulley system and the roof (E). (F) Cartoon demonstrating the FPCT setup and working model with the existence of food pellet and camera. (G) Detailed list of the work pieces and parts of the FPCT setup.

**Supplementary figure 3 (Figure S3).**
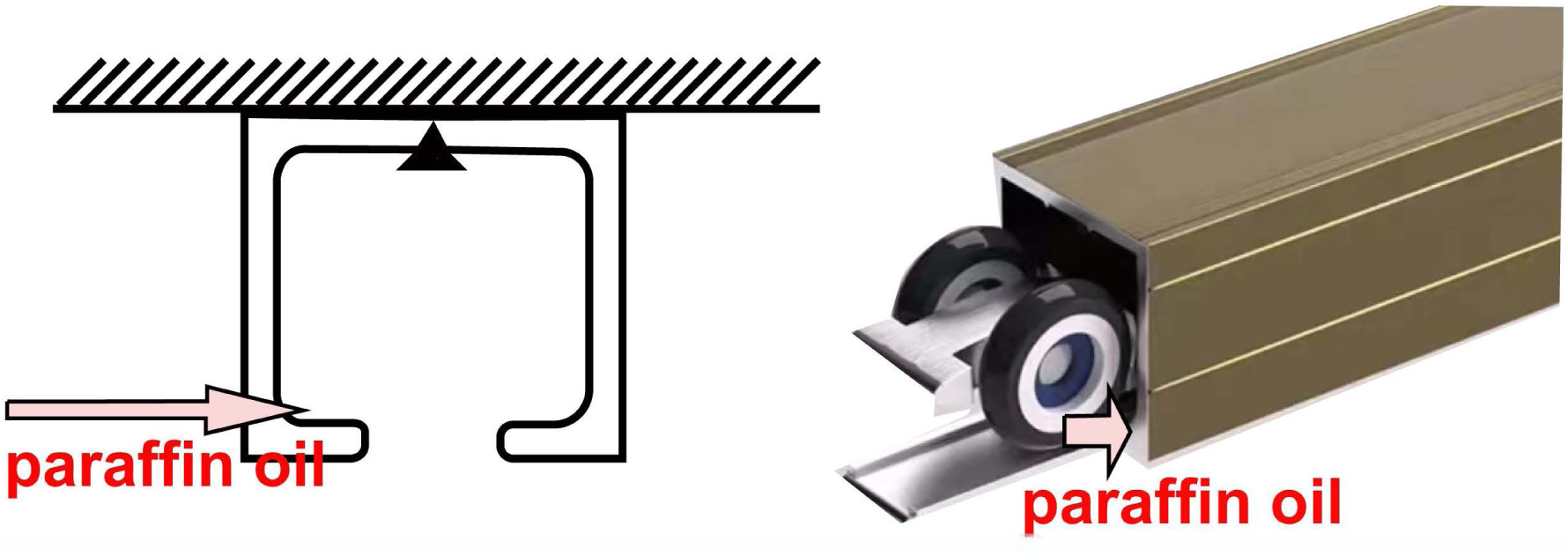
Schematic drawing to show a tip to reduce the friction between the pulley and the track.

**Supplementary figure 4 (Figure S4).**
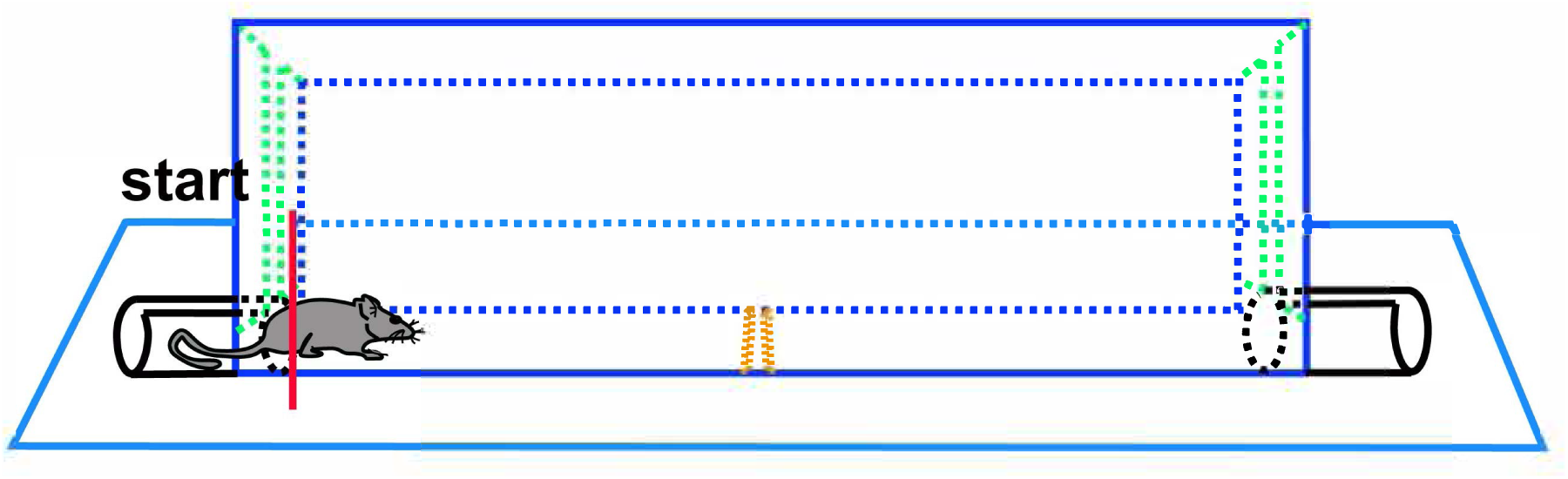
Schematic drawing to show the marking of the starting state when the mouse’s four legs just entered the chamber arena.

## Supplementary videos

**Video S1. In the step 3 of training, the mouse was trained to know the position of the food pellet in the absence of the movable block.**

**Video S2. In the step 4 of training, the mouse was learning to get the food pellet placed in trough of the chamber floor nearly under the movable block.**

**Video S3. In FPCT test 1, the mice were competing for the food pellet.**

**Video S4. In FPCT test 2, the mice were attempting to get the food pellet placed in the inner bottom of the block.**

## Notes

### Competing Interest Statement

The authors have declared no competing interest.

### Summary of Updates

The manuscript has been revised based on the Elife assessment. In response to the reviewers' comments, we 1) re-arranged the figures and provided training data of FPCT; 2) adjusted the introduction and discussion on the history of food competition test; 3) improved the interpretation of our data; 4) described the justification to design FPCT more clearly; 5), corrected the language errors.

